# Combined Angiographic, Structural and Perfusion Radial Imaging using Arterial Spin Labeling

**DOI:** 10.1101/2025.04.29.651006

**Authors:** Thomas W. Okell, Joseph G. Woods, Mark Chiew

**Author notes:** Please address correspondence to: Prof Thomas Okell, FMRIB Centre, John Radcliffe Hospital, Headley Way, Headington, Oxford, OX3 9DU, UK.

## Abstract

**Purpose:** To develop a non-contrast MRI method for the simultaneous acquisition of time-resolved 3D angiographic, perfusion and multi-contrast T1-weighted structural brain images in a single six-minute acquisition.

**Methods:** The proposed Combined Angiographic, Structural and Perfusion Radial Imaging using Arterial Spin Labeling (CASPRIA) pulse sequence uses pseudocontinuous arterial spin labeling (PCASL) to label inflowing blood, an inversion pulse to provide background suppression and T1-weighted contrast, and a continuous 3D golden ratio spoiled gradient echo readout. Label-control subtraction isolates the blood signal and can be flexibly reconstructed at high/low spatiotemporal resolution for angiography/perfusion imaging. The mean signal retains the static tissue, allowing T1-weighted structural images to be reconstructed at different effective inversion times. CASPRIA was compared with conventional time-of-flight (TOF) angiography, 3D-gradient and spin echo (3D-GRASE) PCASL perfusion imaging and magnetization-prepared rapid gradient echo (MP-RAGE) structural imaging (10 minutes total) in healthy volunteers.

**Results:** CASPRIA gave improved distal vessel visibility and fewer artefacts than TOF angiography, whilst also providing dynamic information, with blood transit time and dispersion maps. CASPRIA perfusion images were comparable to 3D-GRASE data, but without through-slice blurring or artefacts in inferior brain regions. Comparable quantitative cerebral blood flow maps were produced, with CASPRIA being significantly more repeatable. Structural CASPRIA images were comparable to MP-RAGE, but also yielded a range of T1-weighted contrasts and allowed quantitative T1 maps to be estimated.

**Conclusion:** CASPRIA is an efficient single acquisition to provide intrinsically co-registered quantitative information about brain blood flow and structure that has considerable advantages over conventional methods.

## 1 Introduction

The use of angiography to visualize blood flow through the brain-feeding arteries is vital in many cerebrovascular diseases, allowing the assessment of stenoses, occlusions or abnormal vessels, such as arteriovenous malformations. However, the effect of any blood flow disruptions on downstream tissue perfusion is also important to observe, since collateral flow and other homeostatic mechanisms can sometimes compensate for compromised arterial flow. In many clinical and research settings, structural images are also required to assess perfusion abnormalities relative to brain structure, to look for abnormal tissues and co-register data from multiple subjects.

Arterial spin labeling (ASL)^1,2^ is an MRI-based approach capable of generating both time-resolved angiograms and quantitative perfusion images non-invasively and without the use of a contrast agent. Conventionally, angiograms, perfusion maps and structural images are acquired separately. This can be time consuming, particularly if high spatial and/or temporal resolution is required, which can make acquiring all of these modalities within a busy clinical protocol infeasible, leaving the clinician with incomplete information.

A method was previously proposed to partially mitigate this issue: Combined Angiography and Perfusion using Radial Imaging and ASL (CAPRIA)^3,4^. This approach allows 4D time-resolved angiographic and perfusion information to be acquired non-invasively from a single scan and has several advantages: a) it is time efficient, since both angiographic and perfusion images can be reconstructed from the same raw k-space data; b) the use of a golden ratio-based readout allows the spatial and temporal resolution of each reconstruction to be chosen retrospectively; c) it does not suffer from significant signal dropout or distortion artefacts common in ASL perfusion imaging; and d) spatiotemporal correlations can be leveraged to improve image quality in reconstruction.

However, CAPRIA also has some drawbacks: a) structural images have to be acquired separately, increasing the total scan time; b) background suppression of static tissue signal is limited, leading to noise amplification, particularly later in the readout during the lower signal-to-noise ratio (SNR) perfusion phase; c) the use of non-selective excitation pulses means a large field of view (FOV) has to be used in reconstruction, increasing the computational burden, as well as including the ASL labeling plane in the reconstructed image, which adds considerable aliased signal from static tissue perturbations due to the labeling process itself; and d) the resulting images were qualitative only.

In this work we propose an extension of CAPRIA to resolve these issues: Combined Angiographic, Structural and Perfusion Radial Imaging using ASL (CASPRIA). We aim to be able to reconstruct all three modalities from a single six-minute scan, maximizing scan efficiency and ensuring perfect co-registration of all the data. In addition, we present models and post-processing strategies to allow quantitative metrics to be extracted from all three modalities and perform initial comparisons of CASPRIA with time- and resolution-matched conventional angiography and ASL perfusion imaging, with additional T1-weighted structural imaging. This builds upon work previously presented in abstract form^5^.

## 2. Methods

### 2.1 Pulse Sequence Design

A schematic of the CASPRIA pulse sequence is shown in Figure 1. It shares some key features with the original CAPRIA approach, including a pre-saturation module to remove spin-history effects and provide some background suppression, a pseudo-continuous ASL (PCASL)^6^ pulse train to label blood water flowing into the brain, and a continuous 3D golden ratio^7^ spoiled gradient echo readout to allow images to be reconstructed with flexible spatial and temporal resolution. The subtraction of PCASL label and control data isolates the blood signal so, for example, a small number of radial spokes can be grouped to reconstruct angiographic images with high spatial and temporal resolution, but with a high undersampling factor that can be tolerated with this high SNR and sparse signal. Perfusion images can be reconstructed with higher sensitivity by grouping a larger number of readouts (lower temporal resolution) using only the central region of k-space where the sampling is denser, improving the conditioning of the reconstruction.

**Figure 1:**
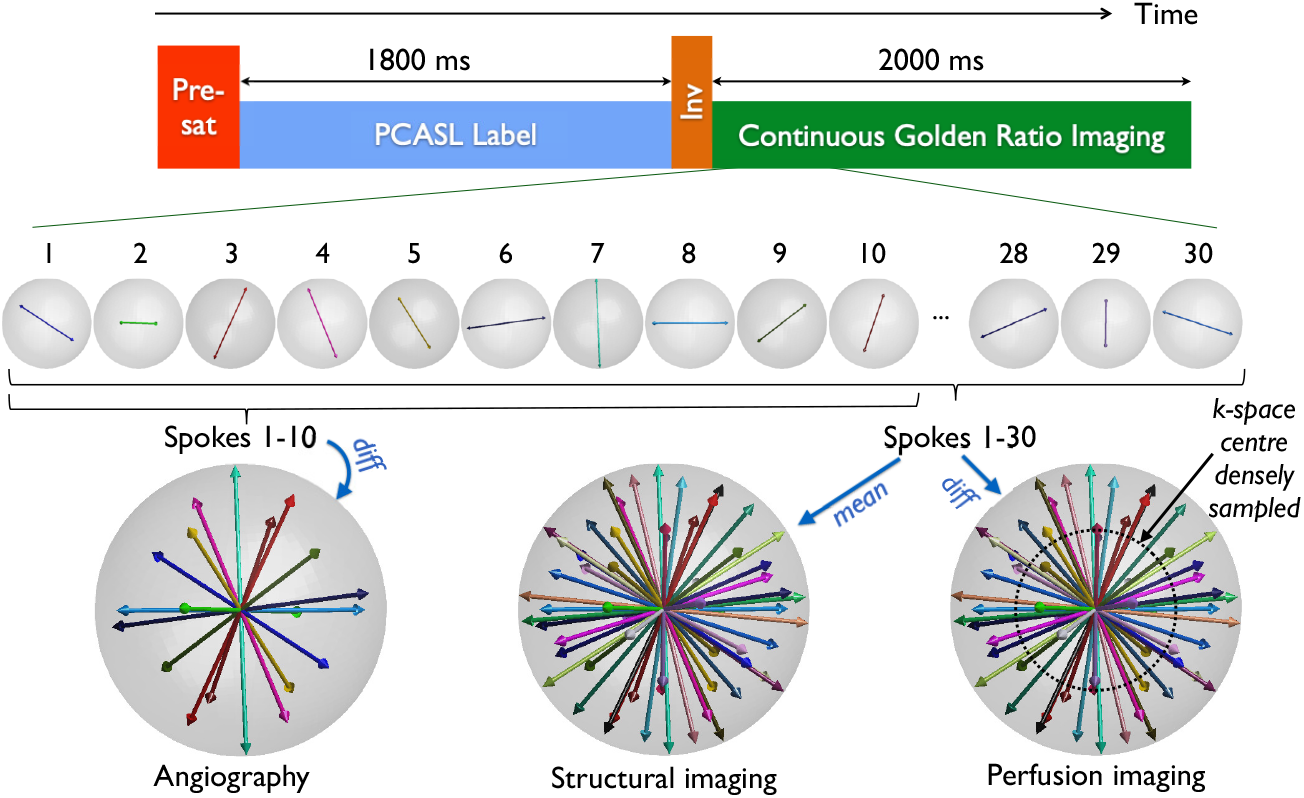
Schematic of the CASPRIA pulse sequence. After a pre-saturation module (“Pre-sat”) and PCASL labeling pulse train, an inversion pulse (“Inv”) is used to reduce the average tissue signal during the long continuous golden ratio 3D radial readout period, as well as providing time-varying T1-weighted contrast. The golden ratio-based readout means that the raw k-space data can be reconstructed at different spatial and temporal resolutions for different purposes: difference data between label and control (“diff”) can be reconstructed at high spatial and temporal resolution for angiography, using a small temporal window, or lower temporal and spatial resolution (using just the more densely sampled centre of k-space) for perfusion imaging, using a larger temporal window. In addition, the mean label/control signal can be taken to retain the static tissue, allowing T1-weighted structural images to be reconstructed at high spatial resolution and low temporal resolution from the same raw dataset. To achieve sufficient k-space sampling, data are combined across many repeats of this process.

However, for CASPRIA, a FOCI inversion pulse (as used in previous studies^8^) has been added immediately after the PCASL pulse train. This both improves background suppression, reducing the average static tissue signal during the readout to better suppress physiological noise, whilst also resulting in time-varying T1-weighted contrast in the static tissue signal, as shown in Figure 2. Rather than only taking the *difference* between label and control data to isolate the blood signal, the *mean* signal can also be used for image reconstruction, retaining the static tissue signal and allowing images with different T1-weightings to be produced at different effective inversion times (TIs) within the long golden ratio readout. With this approach, angiographic, perfusion and structural images can all therefore be reconstructed from a single raw k-space dataset.

**Figure 2:**
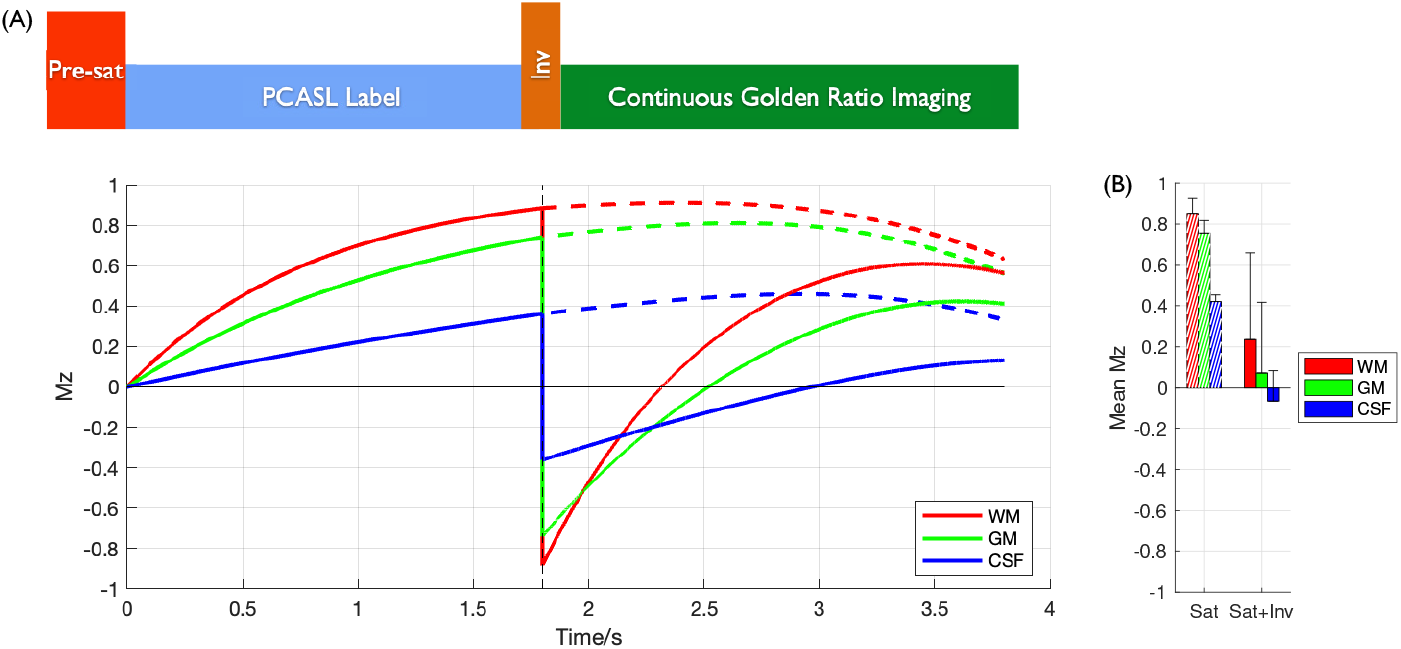
Tissue signal simulations: After the longitudinal magnetization (M_z_) is set to zero by the pre-saturation module, different tissues within the imaging region (white matter [WM], gray matter [GM] and cerebrospinal fluid [CSF], see legend) recover at different rates during the PCASL pulse train (A). In the original CAPRIA implementation, no additional inversion pulse was used (“Sat” only), leading to high tissue signals during the readout period (dashed lines). The additional inversion pulse in CASPRIA (“Sat+Inv”) reduces the average M_z_ during the readout, as shown in (B), but also introduces time-varying T1-weighted contrast during the readout, allowing multiple structural images with different T1-weightings to be reconstructed from the raw k-space data.

A number of additional modifications have been made to allow to improve efficiency and to overcome some of the other drawbacks of CAPRIA mentioned above: a) the PCASL labeling duration has been extended to 1800 ms, to boost the labeled blood signal and bring this approach into line with consensus paper recommendations^9^; b) slab-selective excitation pulses are used during the readout so that a reduced FOV can be reconstructed in the inferior-superior direction, lowering the computational burden and excluding aliased static tissue signal from the labeling plane which can be problematic in these heavily undersampled scans; and c) the radial spoke direction increments according to the golden ratio *first* across repeated ASL preparations and *then* across time, as previously proposed^10,11^, improving the k-space sampling efficiency for any desired temporal resolution.

### 2.2 Subjects and scan protocol

In order to demonstrate the image quality achievable with CASPRIA and compare it to conventional methods, seven healthy volunteers (two female, age range 23 – 44) were scanned under a technical development protocol agreed by local ethics and institutional committees on a 3T Prisma scanner (Siemens Healthineers, Erlangen, Germany) using a 32-channel head coil.

A quick (1min 22s) time-of-flight angiogram was acquired in the neck to enable positioning of the PCASL labeling plane^12^. This was followed by a CASPRIA protocol (~6 mins), and spatial resolution- and total time-matched conventional 3D multi-slab TOF angiography (~3 mins) and 4-shot segmented 3D-GRASE^13^ PCASL perfusion imaging, both multi-postlabeling delay (MPLD) and a single PLD, as suggested by the ASL white paper (WP)^9^, (each 2min 30s + 30s for a calibration (M0) image). In this way, the total time for conventional angiography and one of the perfusion protocols was also approximately 6 mins, matching the CASPRIA scan time. An additional rapid T1-weighted MP-RAGE^14^ scan was also acquired (3min 40s) for comparison with the CASPRIA structural images and an extra 3D-GRASE calibration image (30s) with opposed phase-encoding (left-right rather than right-left) was also acquired to allow for distortion correction of the 3D-GRASE data.

Spatial and temporal resolution were matched as closely as possible between CASPRIA and the comparison protocols, including 1.1mm isotropic resolution for angiography and 3.4mm isotropic resolution for perfusion imaging, with six perfusion postlabeling delays (PLDs) ranging from ~175ms up to ~1800ms (with 1800 ms only for the WP 3D-GRASE protocol). Due to the short time available for the comparison protocols, some compromises had to be made, including a reduced FOV relative to CASPRIA, the use of high parallel imaging (GRAPPA^15^) acceleration factors for TOF angiography and the use of poorer spatial resolution (1.7mm isotropic) for the MP-RAGE. Full details of the protocol parameters are given in Table 1.

**Table 1:**
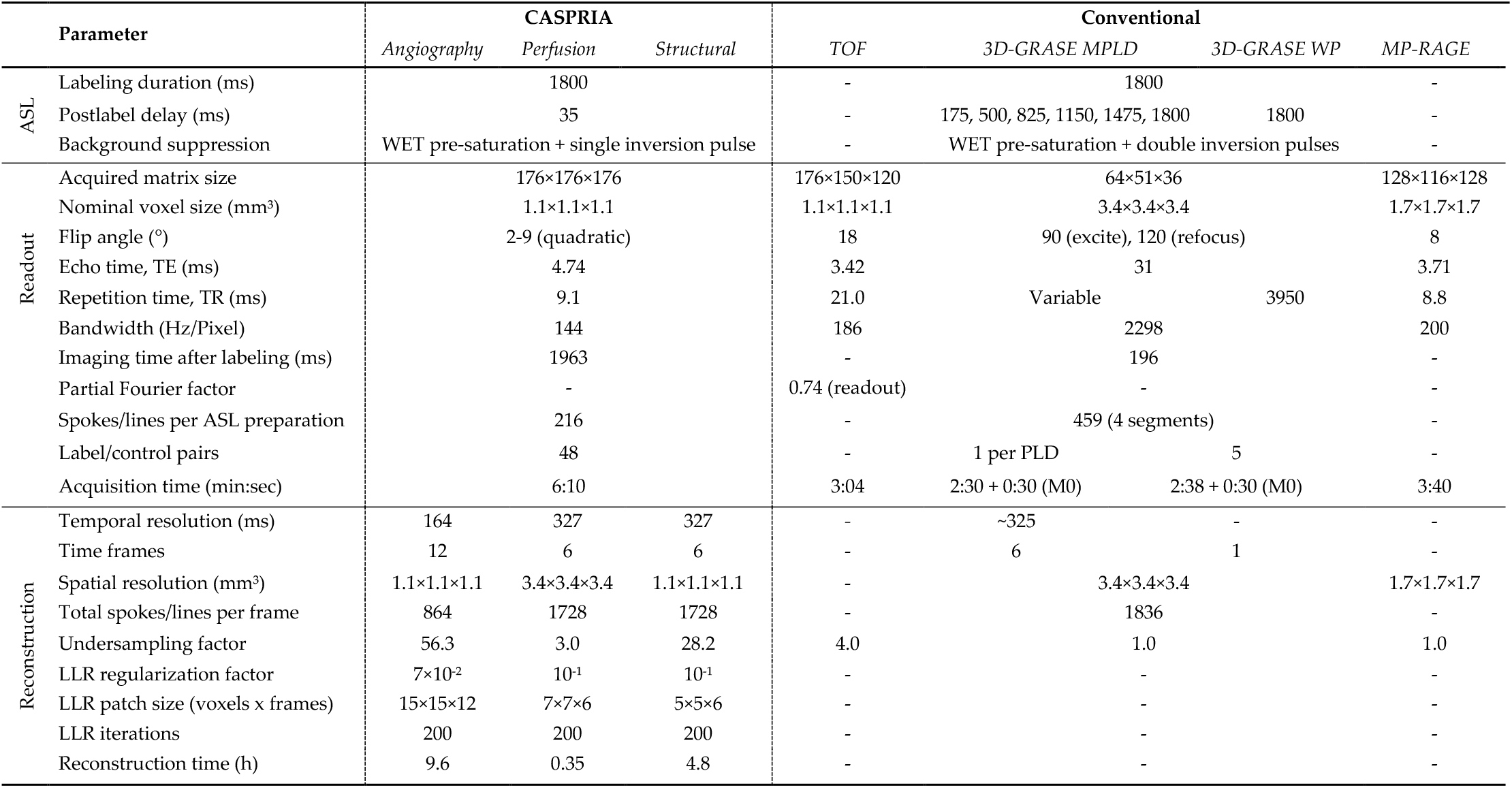
Imaging and reconstruction parameters. Note that only the CASPRIA parameters for protocols B/C is shown here for brevity. Protocol A only differed in the use of 47 label-control pairs rather than 48, with minor changes to scan time (6min 2s vs. 6min 10s) and acceleration factors.

The subjects were scanned with one of three protocols: protocol A (the first 4 subjects) used a CASPRIA scan with 47 ASL label/control pairs (6min 2s) whilst protocols B (subjects 5 and 6) and C (subject 7) used 48 ASL label/control pairs (6min 10s), to evaluate whether the resulting change in the golden ratio ordering affects an artefact seen in preliminary structural CASPRIA reconstructions. Protocols A and B included repeat CASPRIA and 3D-GRASE perfusion scans to allow the assessment of between-scan repeatability, whereas protocol C used only a single repeat to confirm the CASPRIA structural image quality.

### 2.3 Image reconstruction

Locally low rank (LLR) reconstructions are particularly well suited to CAPRIA/CASPRIA data^4^, minimizing signal aliasing and noise amplification whilst improving spatial resolution by leveraging spatiotemporal correlations and the variable k-space sampling patterns across time. This is particularly important for this highly undersampled data, with acceleration factors of 56, 3 and 28 for angiographic, perfusion and structural reconstructions, respectively. The LLR approach^16^, utilizing cycle spinning^17^ and the proximal optimized gradient method^18^, was therefore applied to reconstruct angiographic, perfusion and structural images separately from each raw CASPRIA dataset. Regularization factors and patch sizes were optimized empirically for angiography, perfusion and structural imaging separately, and are given in Table 1.

Coil sensitivities were estimated with the adaptive combine algorithm^19^ and compressed to 8 virtual coils^20^ to reduce the computational burden. Reconstruction was performed on the FMRIB compute cluster (equipped with AMD EPYC 7643 @ 2.3GHz) using four CPUs, taking between ~10 hours for the high spatiotemporal resolution angiographic reconstructions down to ~20 mins for the perfusion reconstructions (see Table 1).

### 2.4 Signal modelling

#### 2.4.1 Angiographic modelling

In order to extract physiological parameters from the CASPRIA dynamic angiograms, we start with the kinetic model for PCASL angiography^21^ and incorporate a modification to account for the variable flip angle (VFA) excitation pulses used during the readout^4^. For this study, since the imaging region is positioned just above the labeling plane, it can be assumed that all of the labeled blood experiences all of the RF excitation pulses, but we keep the effects of dispersion to allow model fitting to real data, leaving us with the angiographic signal:

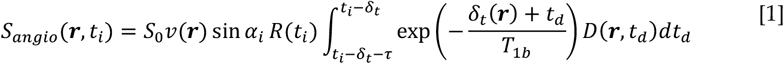

where **r** is the voxel location, t_i_ is the time of the i^th^ RF pulse relative to the start of PCASL labeling, S_0_ is a scaling factor, v is the volume of blood within arteries in that voxel, *α*_*i*_ is the flip angle of the i^th^ RF excitation pulse, *δ*_t_ is the time taken for blood to travel from the labeling plane to the voxel, T_1b_ is the longitudinal relaxation time of blood, t_d_ is an additional time delay due to dispersion, D is the dispersion kernel and R is the additional attenuation of the ASL difference signal due to previous RF pulses, given by:

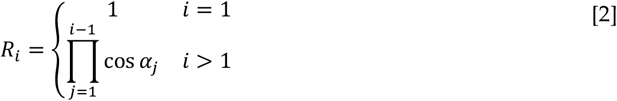

The dispersion kernel, D, has been shown to be well modelled by a gamma variate function^21,22^ using a parameterization similar to Rausch et al.^23^:

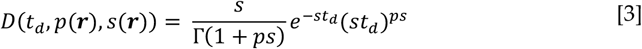

Where Γ is the gamma function, which can be rapidly evaluated numerically:

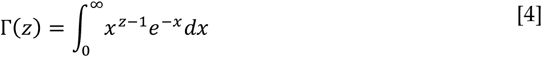

The integral in Eq. [1] can be evaluated numerically^21^, but a faster and more robust solution is possible using the incomplete gamma function, Γ_inc_, which can be rapidly evaluated in many software packages. It can be shown that making the substitution s’ = s + 1/T_1b_ yields a simplified form:

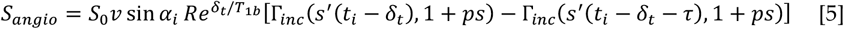

where:

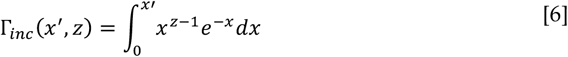

This approach was incorporated in the variational Bayesian model fitting package, FABBER^24^, part of FSL^25^. The modified code can be found at: https://github.com/tomokell/fabber_models_asl/tree/pcasldisp.

This model fitting procedure was applied to CASPRIA angiographic data within a vessel mask defined by taking the temporal maximum intensity projection of the angiogram, multiplying by a brain mask (obtained by applying BET^26^ to a CASPRIA structural image), thresholding at an empirically chosen value to retain most of the vessels, clustering and retaining only the two largest clusters, to remove noisy voxels.

#### 2.4.2 Perfusion modelling

The CASPRIA perfusion signal can be modelled as^4^:

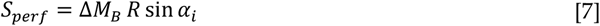

where Δ*M*_*B,i*_ is the Buxton model for the (P)CASL perfusion signal^27^. This modification was also made to the FABBER method, allowing CASPRIA perfusion data to be quantified using the BASIL pipeline^28^, which calls FABBER internally. To compare more directly to the single-PLD (WP) 3D-GRASE data, the final CASPRIA perfusion image at the longest PLD was also isolated and used to quantify CBF on its own.

3D-GRASE perfusion images were processed in a similar manner, with additional inter-volume motion correction and distortion correction using the blip-reversed M0 images via the FSL tool *topup*^29^.

#### 2.4.3 Structural (static tissue) modeling

The static tissue signal changes during the CASPRIA pulse sequence can be derived from simple T_1_ decay and RF attenuation considerations. Here we assume perfect spoiling and a short enough TE to neglect T_2_ decay. After the pre-saturation module, we assume perfect saturation, followed by T_1_ recovery during the PCASL pulse train until just before the centre of the inversion pulse at time 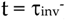, which inverts the magnetization just after the inversion pulse at 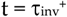 with efficiency *α*_inv_:

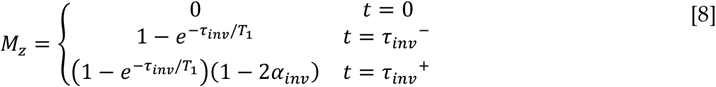

Where here the equilibrium magnetization, M_0_, is set to be 1 for brevity. Further T_1_ recovery occurs until just before the first excitation pulse at t = t_0_:

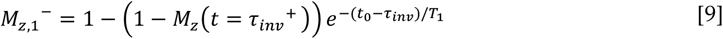

Then the transverse magnetization, M_xy,i_, the longitudinal magnetization just after the RF pulse, 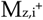, and the longitudinal magnetization just before the next excitation pulse 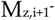 at the i^th^ TR period can be calculated iteratively as:

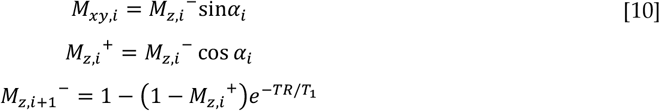

The calculated CASPRIA signal at each timepoint, M_xy,i_, can then be averaged over the time period of each structural image frame and scaled by an additional factor of M_0_.

To fit this model to the CASPRIA structural images, the data were first phase corrected by multiplying through by exp(*-iϕ*), where *ϕ* is the phase in each voxel of the last frame, when the magnetization has recovered to a positive M_z_ state, before the real part of the signal is taken. The MATLAB fitting routine *fmincon* (Mathworks, Natick, MA) was then used to fit for M_0_, *α*_inv_, T_1_ and a relative 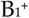 scaling factor, 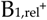 (to account for transmit inhomogeneity), in each voxel by minimizing the sum of squared differences between the data and the model fit. Parameters were constrained to the ranges M_0_ ∈ [0, 10^10^], α_inv_ ∈ [0.5, 1], T_1_ ∈ [0.1, 10] s, 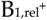 ∈ [0.9, 1.1], and initialized with fits to low spatial resolution (3.4 mm isotropic) data first, which was found to produce more robust results in preliminary testing.

### 2.5 Perfusion calibration procedure

In order to obtain perfusion measurements in absolute units, a measure of the equilibrium magnetization of blood, M_0b_, is required. In conventional PCASL imaging, this is normally achieved by acquiring a separate scan without ASL labeling or background suppression using a long TR to ensure the magnetization is close to equilibrium. For CASPRIA, however, all of the data acquired over the whole scan are used to reconstruct the final image, so producing a calibration scan in the same manner would take as long as the CASPRIA acquisition itself. To avoid the need for any additional scan time, we propose to calibrate the signal using the same raw dataset, as follows.

CASPRIA structural images are reconstructed at the same spatial resolution as the perfusion data (3.4mm isotropic). FSL’s FAST^30^ is applied to the temporal average data to estimate and correct for the bias field. A white matter partial volume map derived from applying FSL’s *fsl_anat* to the MP-RAGE image is linearly registered to the CASPRIA structural images and thresholded at 0.9 to generate a white matter mask. The low resolution CASPRIA structural images are then phase corrected and the real part taken (as described in section 2.4.3) before the mean signal in the white matter mask is taken. The CASPRIA structural model described above is then fit to this curve to extract an estimate of the equilibrium magnetization of white matter. This is converted into an estimate of M_0b_ by dividing by the partition coefficient for white matter (assumed to be 0.82 here).

The CASPRIA perfusion data are then prepared for model fitting by phase correcting (using the low resolution structural images as a reference), taking the real part, multiplying by −1 (to accommodate the single inversion pulse) and then dividing through by the bias field, M_0b_ and the PCASL inversion efficiency (assumed to be 0.85).

Two calibration approaches were used for the 3D-GRASE data: the MPLD data were bias field corrected (using the ratio of M0 images with and without the vendor provided “pre-scan normalize” option) and then a white matter ROI used as the reference region for the BASIL pipeline, to mimic as closely as possible the CASPRIA calibration approach. For the single PLD 3D-GRASE data, the white paper^9^ recommended voxelwise calibration approach was taken.

### 2.6 Image quality assessment

Since the conventional angiography (TOF) and structural imaging (MP-RAGE) provide qualitative data and spatial resolution was lower than CASPRIA for the structural imaging modality, only qualitative comparisons were performed. For the quantitative perfusion data, mean gray matter (GM) perfusion was extracted and scan-rescan repeatability was calculated as the spatial correlation (R^2^) between the repeated scans using cerebral blood flow maps linearly registered^31^ to the MP-RAGE image to ensure alignment. Differences in mean GM CBF and repeatability across the four methods (multi-PLD and single-PLD CASPRIA and multi-PLD and single-PLD 3D-GRASE) were analyzed using a multi-way ANOVA, with imaging method and subject as factors. Where significant effects were found, post-hoc paired t-tests were used to assess differences between pairs of methods.

## 3 Results

Figure 3 shows example 4D angiographic, perfusion and structural images reconstructed from a single raw CASPRIA dataset acquired in six minutes. All frames of these images can be viewed in Supporting Information Videos S1, S2 and S3. Good image quality is observed in all three modalities, clearly showing the blood flowing through the vasculature and perfusing the tissue, with good delineations of brain anatomy and different tissue types with multiple contrasts. Comparison of data acquired protocols A and B/C shows care needs to be taken in choosing the number of ASL preparations in the CASPRIA protocol to avoid structural artefacts due to a complex interaction between spoiling and the golden ratio ordering approach, although angiographic and perfusion images appear to be insensitive to this choice (see Supporting Figures S1 and S2).

**Figure 3:**
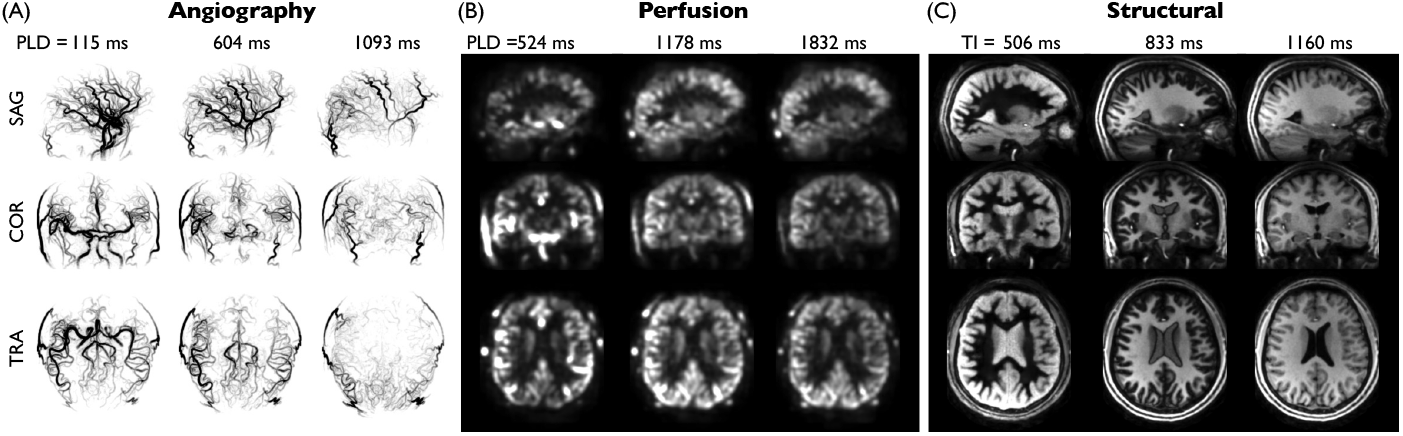
Example angiography, perfusion and structural images reconstructed from a single raw CASPRIA k-space dataset (protocol B) acquired in six minutes: A) Inverted contrast maximum intensity projections (MIPs) of three selected angiographic frames in sagittal (SAG), coronal (COR) and transverse (TRA) views reconstructed with high spatiotemporal resolution, showing the flow of labeled blood through the vascular tree. Note that the use of a long PCASL labeling duration means that the first frame shows much of the vasculature filled with labeled blood. B) Example slices and frames from perfusion images reconstructed at lower spatiotemporal resolution to improve sensitivity, demonstrating the expected patterns of macrovascular signal at early postlabeling delays (PLDs), followed by exchange of labeled blood water into the tissue at later PLDs. C) T1-weighted structural images reconstructed from the average of label and control data at selected times after the inversion pulse (TIs) at high spatial and low temporal resolution, demonstrating the expected contrast changes, including nulling of certain tissues at particular TIs. All frames can be viewed in Supporting Information Videos S1, S2 and S3.

CASPRIA angiography has a number of advantages over the rapidly acquired conventional TOF protocol (Figure 4), including much clearer delineation of distal vessels, lack of venous contamination and slab boundary artefacts, although some small residual loss of signal is observed in some proximal vessels, perhaps due to flow- and/or B_0_-induced dephasing effects. An additional advantage of CASPRIA is the dynamic nature of the data, allowing of visualization of flow through the vascular tree and fitting of a physiological model, as shown in Figure 5. Good fits to Eq. [5] were observed, yielding parameter maps with the expected increases in both blood transit time and dispersion (longer time to peak, *p*, and lower sharpness, *s*) moving towards more distal vessels.

**Figure 4:**
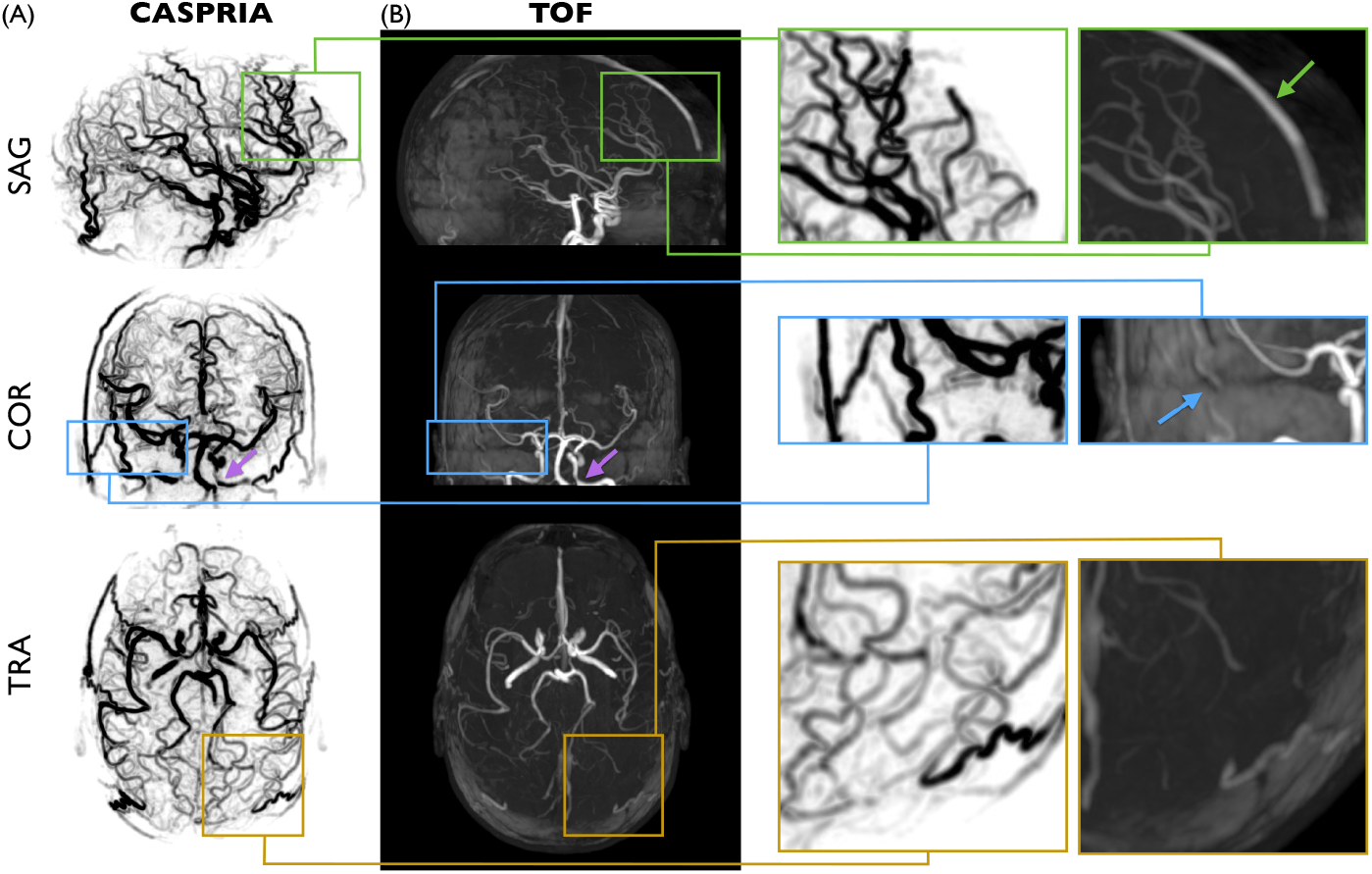
Comparison of CASPRIA angiography with conventional TOF angiography: A) CASPRIA temporal maximum intensity projection across the first seven angiographic frames, shown in different views. B) TOF maximum intensity projections after correction for the non-uniform coil receive sensitivity to reduce overlying tissue interference as far as possible. Zoomed sections and arrows demonstrate the lack of venous contamination signal in CASPRIA that is present in TOF (green), TOF slab boundary artefacts which interrupt vessel signal in slower flowing regions (blue) and the greatly improved visibility of smaller, distal vessels in CASPRIA (yellow). There is some apparent signal reduction in proximal vessels (purple arrows) which is present in both TOF and CASPRIA images, perhaps due to flow- and/or B_0_-based dephasing.

**Figure 5:**
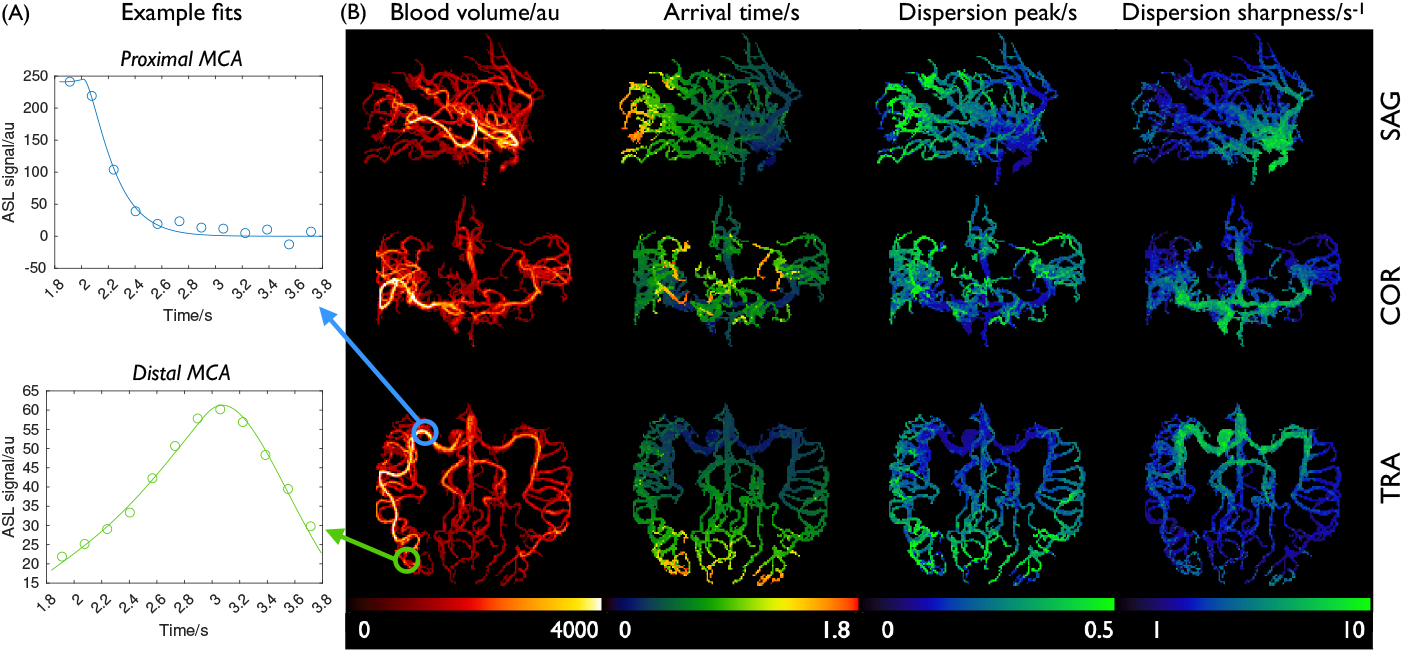
Angiographic model fitting results: A) example fits of the dynamic CASPRIA data to the kinetic model in voxels within the proximal (blue, top) and distal (green, bottom) middle cerebral artery (MCA). B) Maximum intensity projections of the resulting parameter maps, showing the locations of the voxels used in (A) and demonstrating the expected patterns of longer blood arrival time and increased dispersion (delayed peak and lower sharpness) in distal vessels. In these maps the scaling factor S_0_ is merged with the blood volume and estimated as a single scaling factor in arbitrary units.

Dynamic perfusion images generated with CASPRIA also bear a close resemblance to multi-PLD 3D-GRASE perfusion images, as shown in Figure 6, with the expected patterns of macrovascular signal at early PLDs followed by the labeled blood exchanging into the tissue at later PLDs. The macrovascular signal is less apparent in the 3D-GRASE data, most likely due to the use of flow-crushing refocusing pulses. Some of the artefacts seen in the 3D-GRASE data, including through-slice blurring and signal instability in inferior regions, do not appear in the CASPRIA images, which also have higher apparent SNR.

**Figure 6:**
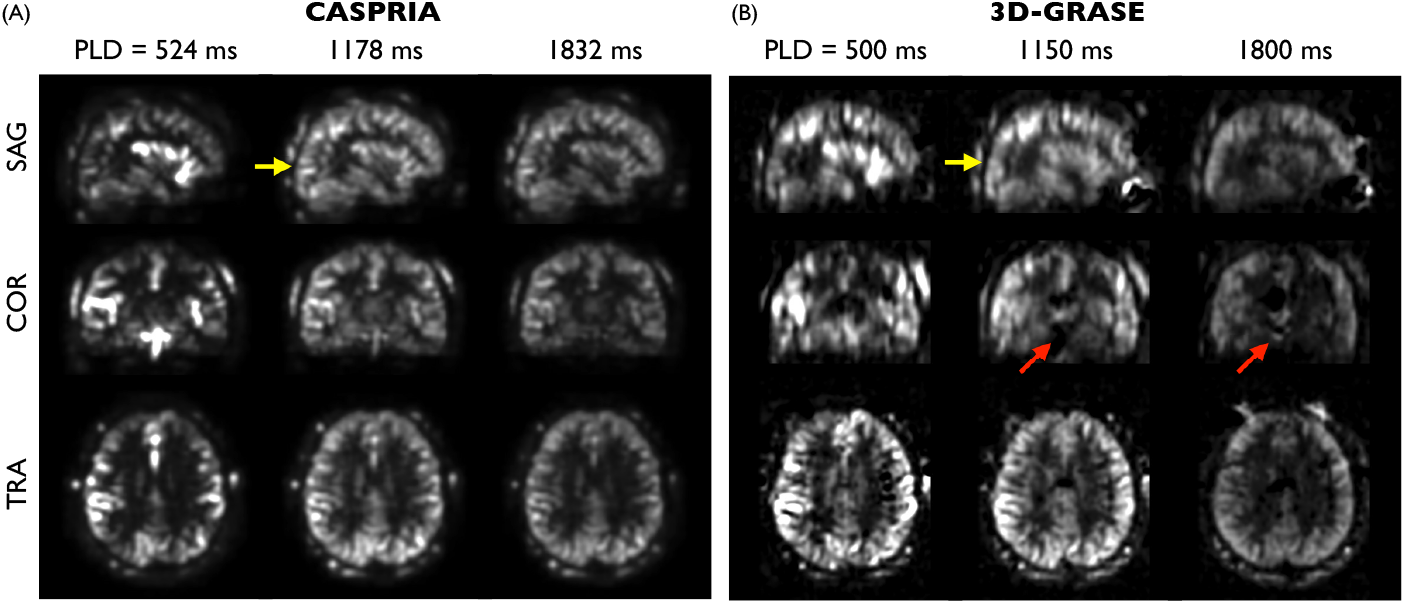
Qualitative comparison of CASPRIA perfusion images (A) with conventional 3D-GRASE multi-postlabeling delay data (B) at selected delay times. Comparable patterns of labeled blood flowing from the vessels into the tissue are seen in both datasets, although the macrovascular signal is more prominent in the CASPRIA data due to the lack of flow-crushing spin-echo refocusing pulses. 3D-GRASE data appears to have lower SNR, increased through-slice blurring (yellow arrows) and some additional artefact in inferior regions (red arrows).

Figure 7 shows the results of the quantitative perfusion comparison. The proposed CASPRIA calibration process appeared to work well, giving good fits to the white matter average signal which allowed CASPRIA perfusion maps to be produced in absolute units with the expected average grey matter values (around 60 ml/100g/min). Absolute CBF maps showed similar features between the CASPRIA and 3D-GRASE approaches, although some residual artefacts were still visible in the 3D-GRASE data and there were absolute scaling differences depending on the method used for calibration, as has been noted previously^32^. Some residual macrovascular contamination may be present in the multi-PLD CASPRIA maps due to the lack of flow-crushing spin-echo pulses during the readout. This was less apparent in the single-PLD (white paper) CASPRIA CBF maps, which were comparable to the 3D-GRASE single-PLD data, with no significant difference in the average grey matter CBF values. In addition, both CASPRIA approaches were significantly more repeatable than both the 3D-GRASE methods, a marker of the improved signal stability.

**Figure 7:**
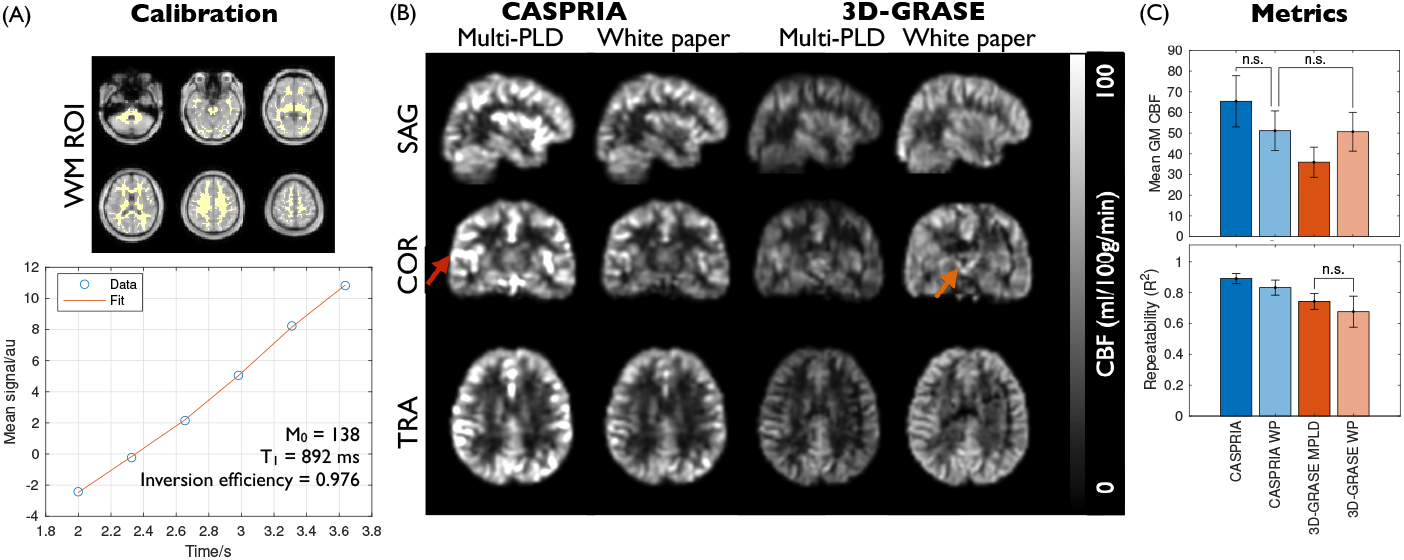
Quantitative perfusion comparison: A) Example CASPRIA calibration procedure using a white matter region of interest (WM ROI). The phase-corrected ROI average signal was fit to the static tissue model to extract estimates of the inversion pulse efficiency, tissue T1 and equilibrium magnetization (M_0_). B) Comparison of quantitative cerebral blood flow maps between CASPRIA and 3D-GRASE (both multi-postlabeling delay [MPLD] and single-delay, white paper [WP] methods). Comparable features are seen in these maps, although there may be some residual macrovascular contamination in the MPLD CASPRIA data (red arrow) and there are global CBF scaling differences. Some residual artefacts are visible in the 3D-GRASE data (orange arrow). C) Mean gray matter CBF metrics (top) reflect the observed global CBF differences across methods, but shows that single PLD CASPRIA gives comparable CBF estimates to single PLD 3D-GRASE. Repeatability metrics (bottom) demonstrate that both CASPRIA approaches are significantly more repeatable than both 3D-GRASE approaches. All differences are significant at p < 0.05 except those marked with “n.s.”.

Finally, example CASPRIA structural images from protocol B are compared with conventional T1-weighted MP-RAGE imaging in Figure 8. The inability to match voxel sizes given the limited imaging time available for the MP-RAGE protocol means a precise comparison is difficult, but comparable tissue contrast can be seen at later CASPRIA TIs. In addition, a variety of other T1-weighted contrasts are produced, including those with some tissues close to their null time, which display even clearer boundaries between tissue types. The multi-TI CASPRIA data can also be used to fit a quantitative T1 map, yielding T1 values in the expected ranges for gray/white matter and cerebrospinal fluid.

**Figure 8:**
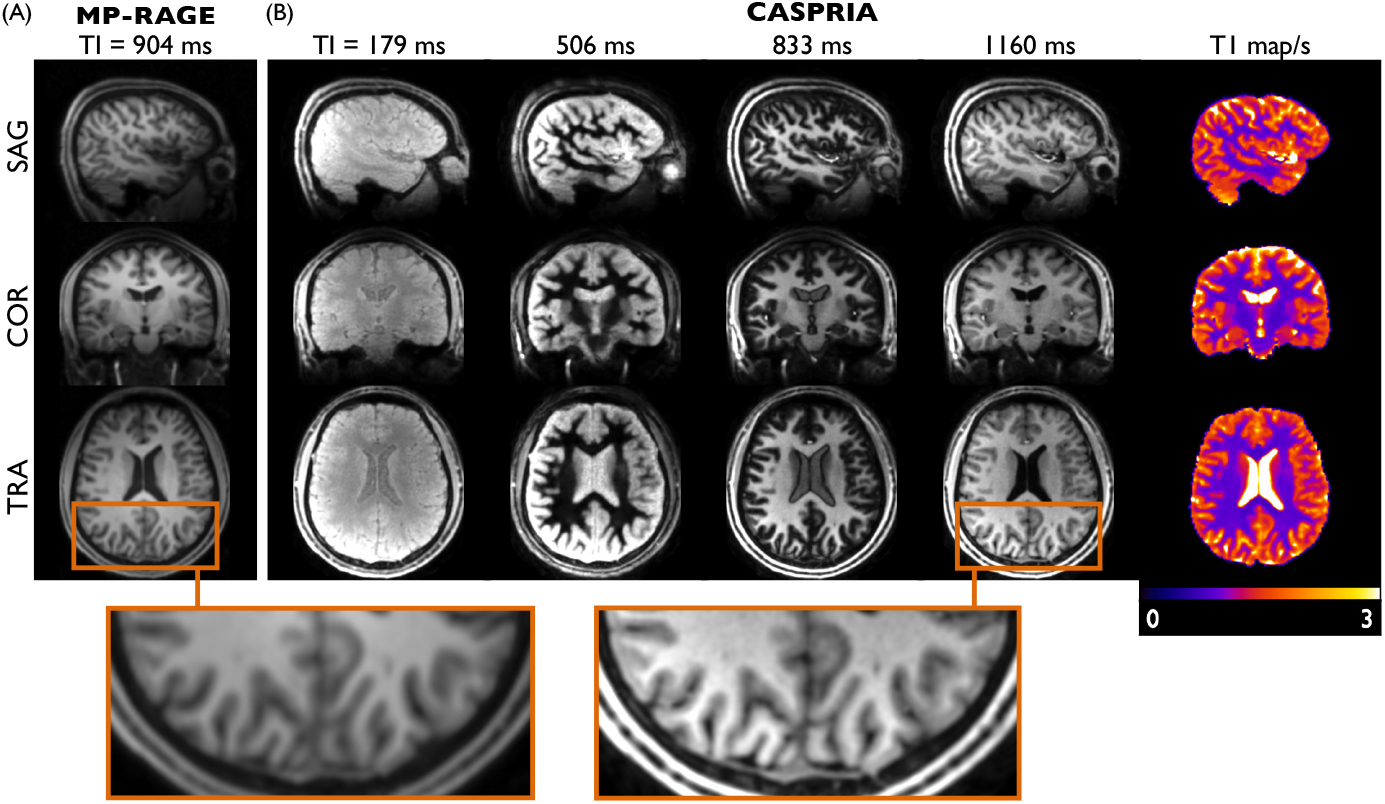
Comparison of conventional MP-RAGE structural imaging (A) with multi-contrast T1-weighted CASPRIA images (B). CASPRIA structural images acquired at later inversion times (e.g. TI = 1160 ms) show comparable tissue contrast and image details to conventional MP-RAGE (see zoomed sections). However, CASPRIA also provides a range of other structural images, including those with certain tissues nulled (e.g. white matter at TI = 506 ms). From this data, a quantitative T1 map can be derived (right column).

## 4 Discussion

The proposed CASPRIA approach generates 4D angiograms, perfusion images and multi-contrast T1-weighted structural images from a single six-minute scan that have significant advantages over conventional methods, which took considerably longer to acquire (~10 minutes in total). CASPRIA angiograms show much clearer depiction of distal vessels, do not suffer from venous contamination or slab-boundary artefacts and are time-resolved, giving additional hemodynamic information (e.g. for separating arterial and venous flow through an arteriovenous malformation) and allowing a kinetic model to be fit to the data, giving additional parameter maps that could be indicative of disease processes^21^. CASPRIA perfusion images give comparable CBF maps to a conventional 3D-GRASE approach but are more repeatable and do not suffer from through-slice blurring or signal instability artefacts. In addition, CASPRIA structural images give comparable contrast to conventional MP-RAGE, but the additional images reconstructed at other TIs give additional information (e.g. with some tissues nulled), which could help with lesion identification, and allow quantitative T1 maps to be estimated.

Similar advantages over conventional TOF for angiography have been noted previously with a similar CAPRIA protocol^4^, although in this study a longer labeling duration and shorter scan time were used, greatly increasing the undersampling factor for CASPRIA, but this did not appear to have a detrimental effect on image quality. In addition, the TOF protocol was better matched in terms of spatial resolution, giving a fairer comparison. The only disadvantage of CASPRIA for angiography is the slight loss of signal in some proximal vessels, most likely due to fast flow and/or B_0_-related dephasing effects, although this could also be partly related to the rapid washout of the trailing edge of the bolus of labeled blood in these proximal vessels that are close to the labeling plane. Improvements could be seen by using a more proximal labeling plane (at a cost of increased T1 signal decay in distal regions) and/or the use of a readout with a shorter echo time, such as the 3D-cones approach^33,34^, which has been recently demonstrated to reduce proximal signal loss for CAPRIA^35^ and will be further explored for CASPRIA in future work.

Perfusion images produced with CASPRIA do not suffer from through-slice blurring artefacts common in ASL perfusion images using 3D readouts that result from T2-induced signal decay during the long readout train. Even though the readout duration was mitigated in this study using a 4-segment 3D-GRASE acquisition, the blurring effect was still noticeable relative to CASPRIA. In addition, the use of segmented 3D-GRASE readouts can lead to signal instabilities, particularly in inferior regions, likely arising from pulsatile motion or respiratory-induced B_0_ field changes.

One notable difference between the modalities is the reduced macrovascular signal seen in 3D-GRASE due to the use of spin-echo refocusing pulses with larger crusher gradients on either side, which dephase magnetization in voxels with a range of flow velocities. This is likely one reason for the differences seen in quantitative CBF maps, particularly for the multi-PLD CASPRIA data, so some further refinement in modelling dispersed macrovascular signal within the perfusion image timeseries is required in the future to mitigate this effect, perhaps leveraging the high SNR angiographic CASPRIA images which are intrinsically co-registered to use as priors in the fitting process. Some global variation in the absolute CBF maps was also noted, most likely due to differences in the calibration procedures, which has been noted previously^32^. This was particularly apparent in the lower CBF values obtained with the multi-PLD 3D-GRASE protocol, perhaps due to the combined use of a scanner receive coil sensitivity correction that does not accommodate B_1_^+^ effects and the use of a white matter reference region, which is mostly concentrated towards the centre of the head, leading to an overestimation of M_0,WM_ and therefore an underestimation of CBF. However, this warrants further investigation in future work.

Using only the final PLD CASPRIA data, however, led to CBF maps with comparable mean gray matter CBF values to the single-PLD white paper 3D-GRASE protocol, showing that the proposed CASPRIA calibration approach was working well. In addition, CASPRIA was shown to be more repeatable than 3D-GRASE, demonstrating that the increased imaging time available through efficiently acquiring multiple modalities simultaneously, as well as sampling the signal over a long readout and potential increases in signal stability arising from the frequent sampling of central k-space, outweighed the lower signal available due to the use of smaller flip angles.

The ability to obtain perfectly co-registered T1-weighted structural images from the same acquisition is a considerable advantage of CASPRIA, and uses similar principles to the previously proposed MPnRAGE method^36^. The golden ratio imaging approach allows for retrospective selection of the TI, which could accommodate images with specific tissues nulled, for example, that could enhance lesion detection or tissue segmentation, although further work is required to determine the optimal post-processing strategies to best leverage this additional information and compare with resolution-matched MP-RAGE data, which was not possible in this study due to scan time constraints. In addition, it was demonstrated that quantitative estimates of tissue T1 can be obtained from fitting the multi-TI CASPRIA data, although comparison with established T1 quantification methods needs to be performed in future work.

This study also highlighted the need to carefully select the number of ASL preparations used in the protocol when using the golden ratio ordering method proposed by Song et al.^10^ and spoiling is performed along the readout direction to avoid accidental refocusing of magnetization from previous TRs. This did not appear to adversely affect the angiographic or perfusion reconstructions, perhaps because the angiographic signal is more rapidly dephased through flow-related effects, making it less prominent in subsequent TRs, and the perfusion images used only the central portion of k-space, away from the observed regions of refocused signal (Supporting Information Figure S2). Another strategy would be to rewind all gradient moments to zero and then use a spoiler gradient in a consistent direction at the end of each TR, although this would require additional time and reduce scan efficiency.

This study has a number of limitations. The use of a single inversion pulse provides useful T1-weighted contrast for structural images, but gives limited background suppression later in the readout, potentially leading to increased physiological noise. Additional inversion pulses during the readout or the use of time-encoded preparations^37,38^ to achieve a shorter readout could improve this, although structural image contrast may be compromised. In this study we used only the LLR reconstruction approach and empirically optimized the reconstruction parameters, which were found to work well on the subjects investigated here, but may need more careful tuning in other cohorts or with other protocols. Only healthy volunteers were investigated, but reoptimization of protocol parameters (e.g. the flip angle schedule) may be required for elderly or patient cohorts. Whilst the 3D golden ratio readout was found to work well here, a similar strategy could be used with other readout schemes, including balanced steady-state free precession^39–41^ readouts, or alternative trajectories such as stack-of-stars^40^, multi-slice golden angle radial methods^11^ or 3D cones^35^. Further improvements are also possible through recent developments such as subspace reconstructions^42,43^ and motion correction^44^, which will be explored in future work.

## 5 Conclusions

The proposed CASPRIA approach allows the generation of 4D angiographic, perfusion and structural images in a single six-minute scan, with significant advantages in vessel-visibility, reduced artefacts, increased signal stability and time-resolved information over separately acquired conventional acquisitions taking 10 minutes.

## 6 Acknowledgements

We are grateful for funding support from a Sir Henry Dale Fellowship jointly funded by the Wellcome Trust and the Royal Society (220204/Z/20/Z). The Wellcome Centre for Integrative Neuroimaging is supported by core funding from the Wellcome Trust (203139/Z/16/Z) with additional support from the NIHR Oxford Health Biomedical Research Centre (NIHR203316). MC is supported by the Canada Research Chairs program. The views expressed are those of the authors and not necessarily those of the NIHR or the Department of Health and Social Care. Many thanks also to Jeff Fessler, Philipp Ehses and colleagues for making available their excellent NUFFT and Siemens raw data reading MATLAB code, as well as to Siemens Healthineers for providing the base pulse sequence code that we built upon in this work. For the purpose of open access, the author has applied a CC BY public copyright license to any Author Accepted Manuscript version arising from this submission.

## 8 Tables

## 9 Figures

**Supporting Figure S1:**
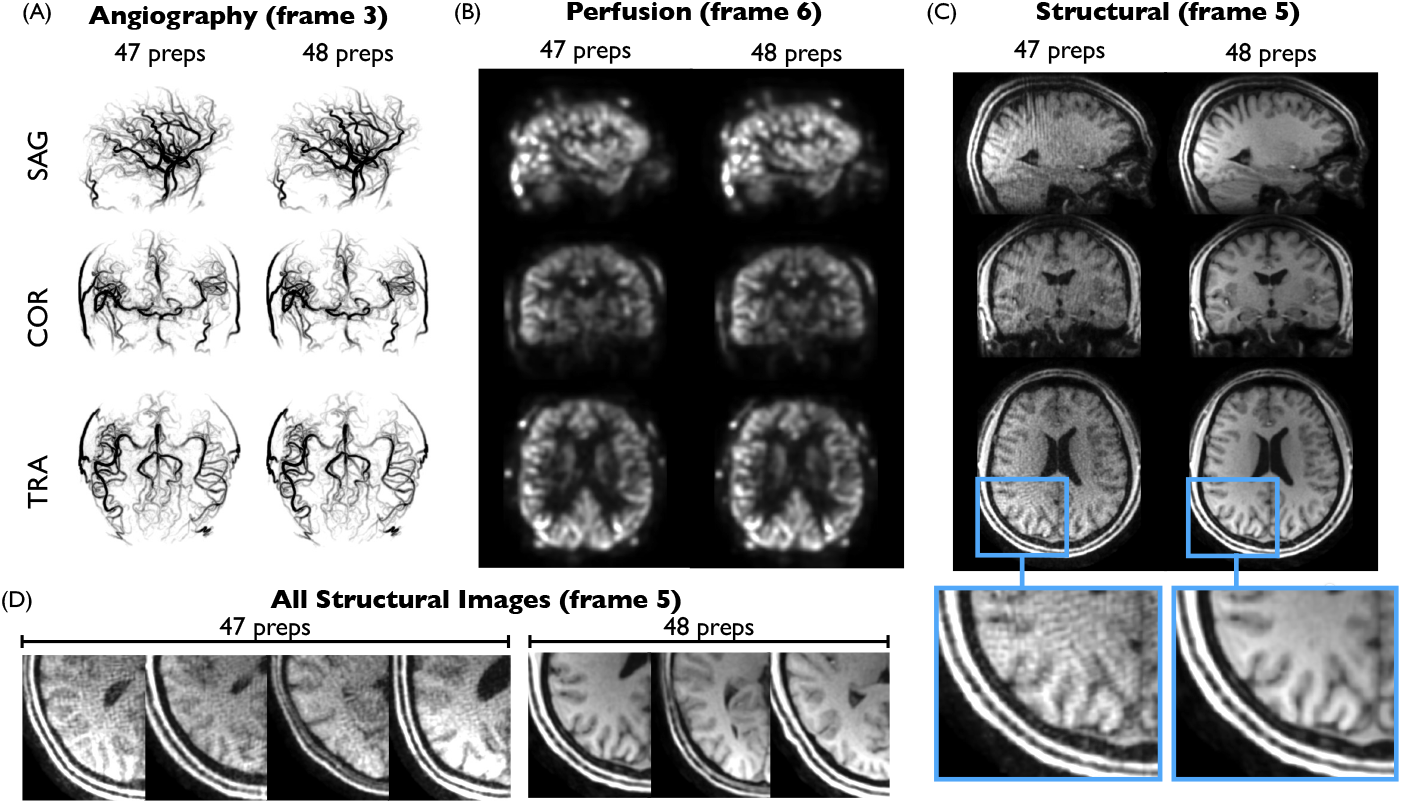
Golden ratio related structural artefacts: example angiographic (A), perfusion (B) and structural (C) CASPRIA images in the same subject acquired with 47 or 48 ASL preparations. Angiographic and perfusion images are almost indistinguishable, with no apparent artefacts, but the CASPRIA structural image reconstructed from the 47 ASL preparation scan has a wave-like patterns (see zoomed section) suggestive of excess signal at higher spatial frequencies. These artefacts are not present in the 48 ASL preparations protocol. This artefact was consistently seen across the four subjects which used the 47 preparations protocol and was not present in all subjects that used the 48 preparations protocol (D), confirming this to be the source of the problem.

**Supporting Figure S2:**
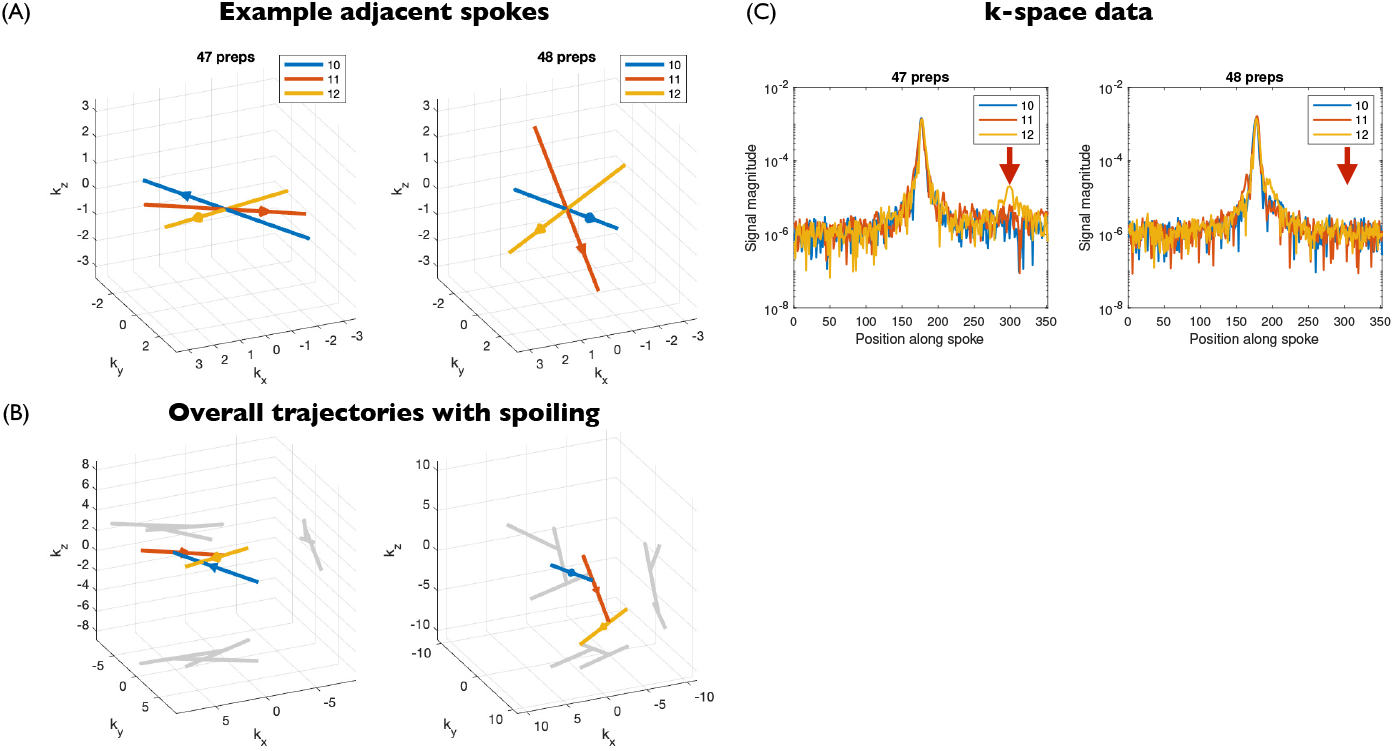
Comparison of k-space trajectories with the two CASPRIA protocols used in this study (with 47 or 48 ASL preparations). The use of the Song et al.^10^ approach to golden ratio looping, which increments spoke angles with the golden ratio *first* down repeats *and then* across time, means that consecutively acquired spokes in time can have many golden ratio increments between them. When 47 ASL preparations are used, this can result in adjacent spokes being close to anti-parallel in some cases (spokes 10-12 acquired after ASL preparation 4 are shown in A). After each spoke, a spoiler gradient is added in the same direction as the readout gradient, but if the total trajectory of all three spokes with spoilers are concatenated (B) it can be seen that magnetization excited by excitation pulse 10 can be largely refocused during the acquisition of spoke 12. In contrast, if 48 ASL preparations are used, the spoke distribution is improved, leading to previously excited magnetization being further dephased during the acquisition of the next spoke. This refocused signal can be seen in the acquired data with 47 ASL preparations (C, red arrow), which is avoided when 48 ASL preparations are used.

**Supporting Information Video S1**: Video of the CASPRIA angiography maximum intensity projections from one subject shown in Figure 3.

**Supporting Information Video S2**: Video of the CASPRIA perfusion images from one subject shown in Figure 3.

**Supporting Information Video S3**: Video of the multi-contrast CASPRIA structural images from one subject shown in Figure 3.

